# Longitudinal trajectories of aperiodic EEG activity in early to middle childhood

**DOI:** 10.1101/2024.09.10.612049

**Authors:** Dashiell D. Sacks, Viviane Valdes, Carol L. Wilkinson, April R. Levin, Charles A. Nelson, Michelle Bosquet Enlow

## Abstract

**Background:** Emerging evidence suggests that aperiodic EEG activity may follow a nonlinear growth trajectory in childhood. However, existing studies are limited by small assessment windows and cross-sectional samples that are unable to fully capture these patterns. The current study aimed to characterize the developmental trajectories of aperiodic activity longitudinally from infancy to middle childhood. We examined potential trajectory differences by sex and brain region. We further investigated whether aperiodic activity is associated with maternal anxiety symptoms, and whether these associations vary because of differential development trajectories.

**Methods:** A community sample of children and their parents (*N*=391) enrolled in a longitudinal study of emotion processing were assessed at infancy, and at ages 3 years, 5 years, and 7 years. Analyses included individual growth curve and mixed effect models. Developmental trajectories of the aperiodic slope and offset were investigated across whole brain, frontal, central, temporal, and posterior regions. Associations of whole brain slope and offset with maternal anxiety symptoms were also examined.

**Results:** Developmental trajectories for both slope and offset were generally characterized by a relative increase in early childhood and a subsequent decrease or stabilization by age 7, with variation by brain region. Females showed relatively steeper slopes at some ages, and males showed relatively greater offset at certain ages. Maternal anxiety was negatively associated with slope at 3 years and positively associated with slope at 7 years.

**Conclusions:** The longitudinal developmental trajectory of aperiodic slope in early childhood is nonlinear and shows variation by sex and brain region. The magnitude and direction of associations with maternal anxiety varied by age, corresponding with changes in trajectories. Developmental stage should be considered when interpreting findings related to aperiodic activity in childhood.

---

A large body of research has focused on delineating the properties and functions of electroencephalography (EEG) power spectral density, with the goal of elucidating biological indicators (biomarkers) of brain functioning. EEG is portable and affordable relative to other neuroimaging methods. Thus, EEG is an appealing modality for investigating such biomarkers, which may facilitate early identification and intervention strategies for improving developmental outcomes. Traditional analytical methods typically involve averaging periodic oscillatory power within putative, predefined frequency bands, including delta, theta, alpha, beta, and gamma. However, the EEG power spectrum also contains an aperiodic component, which is characterized by a variable 1/f^x^ distribution in which power decreases as frequency increases. This aperiodic activity has traditionally been viewed as “background noise.” Consequently, canonical analytical methods tend to conflate aperiodic activity with periodic activity, or actively attempt to reduce or eliminate aperiodic activity from analyses (Donoghue et al., 2020; Levin et al., 2020). Recent methodological advances, particularly spectral parameterization (SpecParam; previously fitting oscillations of one/f) enable parameterization of the power spectrum into distinct variables that represent both periodic and aperiodic components.

The aperiodic component reflects non-oscillatory neuronal spiking activity (Gao et al., 2017; Manning et al., 2009; Miller et al., 2009) and is defined by the aperiodic “offset” and “slope.” The offset indexes the uniform shift in power across EEG frequencies; the slope indexes the rate at which power increases as frequency decreases, defined by the exponent x in the 1/f^x^ distribution. Aperiodic activity, particularly slope, has started to receive attention in various studies, with more recent hypotheses proposing that aperiodic activity reflects mechanistic underpinnings of brain activity (He et al., 2010). Specifically, the slope has been described as the “signal to noise ratio” of neural activity and as a potential index of the balance between synaptic excitation and inhibition (excitatory-inhibitory [E-I] balance; (Donoghue et al., 2020; Gao et al., 2017). According to this conceptualization, flattened or reduced slope is hypothesized to reflect increased excitatory activity over inhibitory activity. In contrast, steeper or more accelerated slope is hypothesized to reflect increased inhibitory activity over excitatory activity. However, further research is required to establish these primarily theoretically driven assumptions.

Cross-sectional studies investigating slope across the lifespan, starting from 4 years of age and through late adulthood, report reduced or “flattened” slope in older participants compared to younger participants (Hill et al., 2022; Merkin et al., 2023; Voytek et al., 2015). However, a recent cross-sequential study by Wilkinson et al. (2024) reported the opposite trend during early childhood from 2 to 44 months of age. Specifically, slope increased or “steepened” during early childhood, particularly during infancy. Furthermore, McSweeney et al. (2023) fit mixed models both with and without an added quadratic term to investigate slope trajectories (cross-sectionally) across 4- to 11-year-olds. They found the model with an added quadratic term provided a significantly better fit, suggesting nonlinear developmental trends throughout this age period. Of the aforementioned studies, Wilkinson et al. (2024) was the only one to test sex effects. Significant age-by-sex interactions were reported, with sex differences emerging in models after approximately the first year of life. Steepening slope in early childhood could be reflective of increases in brain volume and synaptogenesis that occur uniquely during early development. If slope is associated with E-I balance, increasing slope in early childhood may also reflect the development in inhibitory networks that occurs during early childhood. Whereas excitatory neurons are well established by birth, GABAergic (Gamma-Aminobutyric Acid) inhibitory neurons continue to migrate from ventral subregions of the brain to the cortices in early childhood (Chini et al., 2022; Paredes et al., 2016).

The signal-to-noise ratio hypothesis suggests that the association between slope and brain functioning may be monotonic, with steeper slope reflecting an enhanced signal-to-noise ratio. However, if slope indexes E-I balance, the optimal slope may follow a more parabolic or inverted U-shape. Such a pattern would be aligned with the complex systems approach, which suggests that there are nonlinear dynamics in biological systems, as opposed to the linear relationships often observed in more simple systems (Cohen et al., 2022). In line with this framework, there may be an optimal homeostatic range, such that slopes that are either too flat or too steep are associated with cognitive impairment. Robertson et al. (2019) further posit that there may be a range of cognitively optimal slopes at different developmental stages. Differential trajectories of slope may be associated with different optimal slope ranges. Furthermore, cortical maturation occurs in a regionally differential, posterior-anterior pattern (Gerván et al., 2017; Shaw et al., 2008), which suggests the potential for regionally based differences in slope trajectories. Although previous research that characterized linear trajectories of slope reported similar trends across regions (Hill et al., 2022), this research may have been limited in its ability to capture more nuanced differences by taking a linear approach and relying on a cross-sectional design.

If slope is characterized by nonlinear developmental trajectories and dynamics, differential associations with neuropsychiatric outcomes and environmental influences known to affect brain development may be observed at different stages of development. Recent studies at different ages in childhood and adulthood have demonstrated associations between both steeper and flatter slope and various neurodevelopmental and mental health disorders (Karalunas et al., 2022; Robertson et al., 2019). Research is required to investigate whether these differences may relate to developmental dynamics. Early childhood marks a period of substantial vulnerability to environmental influences for brain development. Significant developmental processes, including axonal and dendritic growth, synaptic stabilization, and synaptic pruning are ongoing throughout childhood (Chen & Baram, 2016). Caregivers are one of the most salient environmental influences during early development, with caregiver psychopathology a known risk factor for differences in offspring neurodevelopment (Chen & Baram, 2016). Here we investigated whether any differences in aperiodic activity trajectories by developmental stage influenced the nature of associations between aperiodic activity and maternal anxiety across the first 7 years of life.

## The Current Study

The extant literature suggests there may be a shift in early childhood from increasing (steeper) to decreasing (flattening) slope (Hill et al., 2022; McSweeney et al., 2023; Voytek et al., 2015; Wilkinson et al., 2024). Longitudinal research is required to investigate developmental shifts during childhood. It is important for such investigations to examine the influence of sex on slope trajectories, given reported sex differences in early childhood (Wilkinson et al., 2024). Investigations should also consider potential differences by brain region, given typical regional differentiation in patterns of brain development throughout childhood (Gerván et al., 2017; Shaw et al., 2008). Research is also needed to understand how trajectories of aperiodic EEG may influence the nature of associations with early environmental exposures, such as maternal psychopathology symptoms. The primary objective of the current study was to investigate the developmental trajectories of aperiodic activity across the first 7 years of life in a longitudinal community sample of typically developing children. Specifically, the aims were to: 1) characterize the developmental trajectories of electroencephalography (EEG) power spectral density slope and offset from infancy to middle childhood in whole brain and frontal, central, temporal, and posterior regions; 2) examine potential sex differences in developmental trajectories; and 3) explore whether shifting developmental trajectories from infancy to 7 years influence the nature and magnitude of associations with maternal anxiety, a known influence on child neurodevelopmental outcomes. Aims 1 and 2 were primarily confirmatory, whereas Aim 3 was exploratory.

## Method

### Participants

Participants were recruited at infancy from a registry of local births in the Greater Boston Area, MA, United States of America, comprising families who had indicated willingness to participate in developmental research. Families in the current analyses participated in a prospective longitudinal study to examine the early development of emotion processing. Exclusion criteria included known prenatal or perinatal complications, maternal use of medications during pregnancy that may significantly impact fetal brain development (i.e., anticonvulsants, antipsychotics, opioids), pre- or post-term birth (±3 weeks from due date), developmental delay, uncorrected vision difficulties, and neurological disorder or trauma. After enrollment, families were no longer followed, and their data were excluded from analyses if the child was diagnosed with an autism spectrum disorder, or a genetic or other condition known to influence neurodevelopment. By design, families were enrolled in the parent study when the children were 5, 7, or 12 months old, with a subsample followed at 3 year, 5 years, and 7 years. In the current analyses, *N*=391 children provided data at infancy; of these 391 participants, *N*=187 provided follow-up data at 3 years, *N*=160 at 5 years, and *N*=106 at 7 years. This study builds on Wilkinson et al. (2024), which reported trajectories of aperiodic activity from infancy through age 3 years, by investigating a subset of participants through age 7 years.

### Procedures

Parents (almost exclusively the child’s mother [> 98%]; hereafter referenced as “mother”), were asked to complete questionnaires via an online survey prior to or during each in-person study visit (infancy, 3 years, 5 years, 7 years). Questionnaires relevant to the current analyses included assessments of sociodemographic characteristics collected at a baseline assessment in infancy, and maternal anxiety symptoms collected at all time points. During each in-person study visit, EEG data were collected. The Institutional Review Board at Boston Children’s Hospital approved all study methods and procedures, and parents provided written informed consent prior to the initiation of study activities. Initial infancy study visits began in April 2013 and ended in April 2017. Data collection for the 7-year visits continued through October 2023.

### Measures

#### Sociodemographic Characteristics

Sociodemographic characteristics collected included child age, sex assigned at birth (hereafter “sex”), ethnicity, parental education, and annual household income.

#### Maternal Anxiety Symptoms

Maternal anxiety symptoms were measured at infancy and 3, 5, and 7 years using the Trait Anxiety form of the Spielberger State-Trait Anxiety Inventory (STAI-T; Spielberger, 1983). The STAI is a 20-item self-report questionnaire designed to measure anxiety proneness. Respondents are asked to rate the frequency of general mood states on a 4-point scale, ranging from ‘almost never’ to ‘almost always.’ Item scores were summed to create a total score (possible range: 20 – 80), with a higher score indicating greater anxiety. The STAI-T has been established to have good internal consistency (α=.90), and test-retest coefficients from .73 to .86 (Barnes et al., 2002). Internal consistency estimates for this sample were: α=.89 at infancy, α=.89 at 3 years, α=.91 at 5 years, and α=.91 at 7 years.

#### Electroencephalography (EEG)

##### EEG Acquisition

Continuous scalp EEG was recorded using a 128-channel HydroCel Geodesic Sensor Net (HGSN; Electrical Geodesic Inc.) with Ag/AgCl coated, carbon filled electrodes. The four electrooculogram channels were removed from the net for infant comfort. The net was connected to a NetAmps 300 amplifier (Electrical Geodesic Inc.) and referenced online to a single vertex electrode (Cz). Data were sampled at 500 Hz. Channel impedances were kept at or below 100 kΩ, which is within recommended guidelines given the high-input impedance capabilities of the system’s amplifier. At infancy and 3 years, participants watched a computer-generated video of slow-moving infant toys in a dim room, while baseline EEG (2 minutes) was recorded. Participants sat on their caregiver’s lap for the infancy visit. At 5- and 7-years participants watched a computer-generated screen-saver style video of flashing lights while baseline EEG was recorded (4 minutes).

##### Preprocessing

Raw Net Station (Electrical Geodesics, Inc) EEG files were exported into the MATLAB MAT-file format for preprocessing in MATLAB (version R2023a), using the Batch Automated Processing Platform (BEAPP) (Levin et al., 2018), with integrated Harvard Automated Preprocessing Pipeline for EEG (HAPPE) (Gabard-Durnam et al., 2018). Data were high-pass filtered at a 1Hz and low-pass filtered at 100Hz. Data were resampled to 250Hz, and then preprocessed using the HAPPE module, which includes line noise removal, bad channel rejection, and artifact removal using combined wavelet-enhanced independent component analysis (ICA) and Multiple Artifact Rejection Algorithm (MARA; (Winkler et al., 2015). The following channels were used in addition to the 10-20 electrodes for MARA: 28, 19, 4, 117, 13, 112, 41, 47, 37, 55, 87, 103, 98, 65, 67, 77, 90, 75. After artifact removal, bad channels were interpolated. Data were referenced to the average reference, detrended to the signal mean, and segmented into 2-second epochs. HAPPE’s amplitude and joint probability criteria were used to reject epochs contaminated with artifact. Recordings with fewer than 20 segments (40 seconds of total EEG), percent good channels < 80%, percent independent components rejected >80%, mean artifact probability of components kept > 0.3, or percent variance retained < 25% were rejected. Across the full sample, there were *n* = 100 (12%) data points rejected.

##### Power spectral density

Power spectral density was calculated in the BEAPP Power Spectral Density (PSD) module using a multitaper spectral analysis with three orthogonal tapers. For each electrode, the PSD was averaged across segments, and then further averaged across all channels to derive the whole brain average. PSD was further averaged in electrodes across frontal, central, temporal, and posterior ROIs (Figure 1).

**Figure 1.**
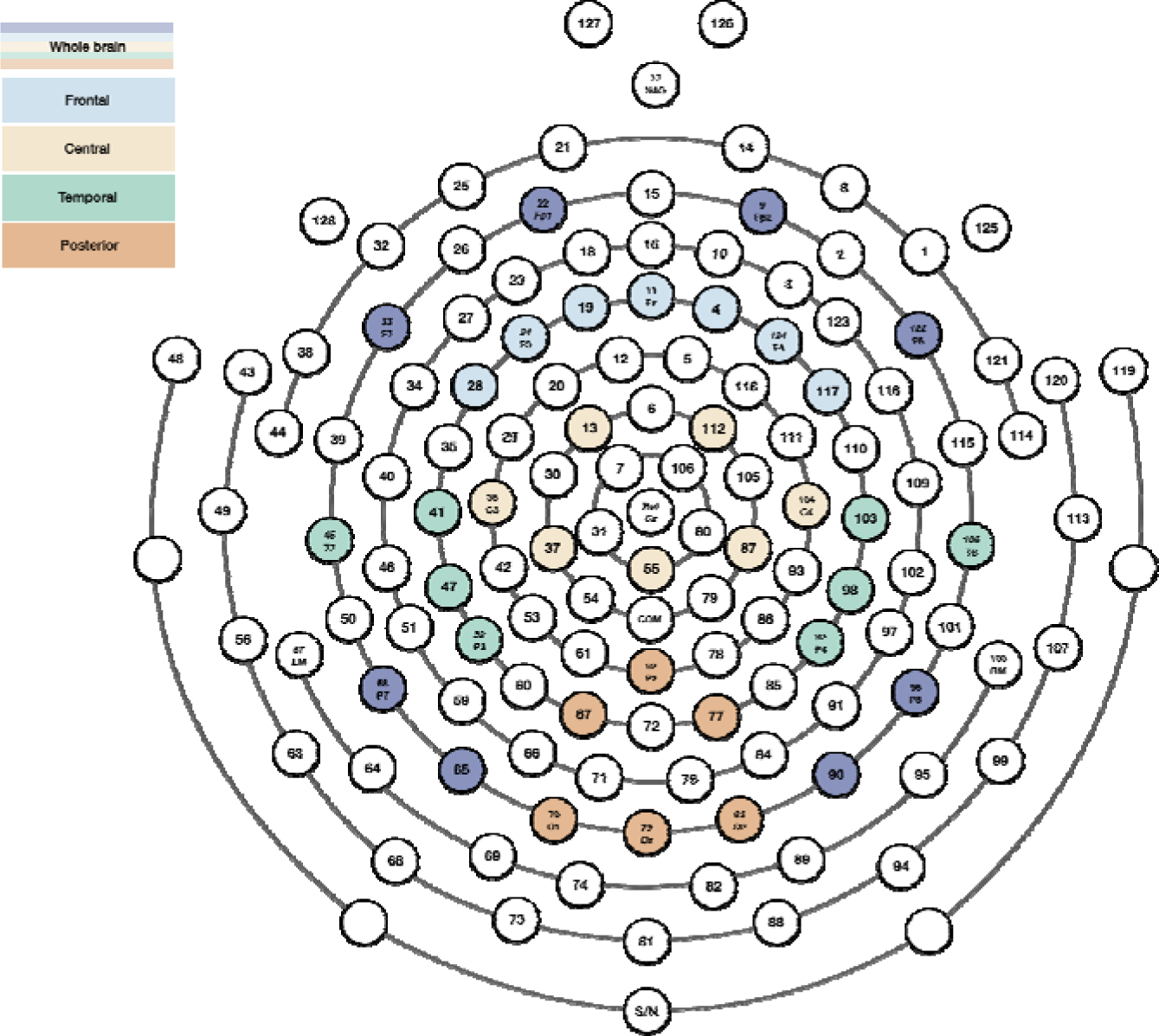
Electrode Layout for Whole Brain and Regions of Interest. Note: Regions of interest are color-coded as frontal (light blue), central (yellow), temporal (green), and posterior (orange). Whole brain channels are derived from a combination of the electrodes used for each region and the electrodes marked in dark blue.

##### Spectral Parametrization

The PSD was parameterized using SpecParam (Donoghue et al., 2020). The adapted version originally developed by Wilkinson et al. (2024) was adopted to improve model fit in the infant EEG. In this version, the robust_ap_fit function is modified, so that the initial estimate of the flattened power spectra (flatspec) has a baseline elevated such that the lowest point is ≥ 0. Further details are available in the papers by Donoghue et al. (2020) and Wilkinson et al. (2024). The modified version had limited impact at later ages. Pearson’s correlation values between whole brain slope calculated using the original and modified scripts were .97, .98, and .99 at 3, 5, and 7 years respectively. Correlation values between whole brain offset calculated using the original and modified scripts were and were .96, .97, and .98 at 3, 5, and 7 years respectively. The SpecParam model was used across a 2–55 Hz frequency range, in the fixed mode (no spectral knee) with peak_width_limits set to [0.5, 18.0], max_n_peaks = 7, and peak_threshold = 2. Mean R^2^ ranged from 0.994 – 0.998 across each age group and region, *SD* ranged from 0.002 – 0.030. Mean estimated error for the sample at across each age group and region ranged from 0.013 – 0.020, *SD* ranged from .006 – 0.016.

### Statistical Analysis

Statistical analyses were conducted using the Statsmodels (Seabold & Perktold, 2010) package in Python 3.10.10. Descriptive statistics for the sample’s sociodemographic characteristics and main study variables were calculated. Slope and offset trajectory analyses were conducted for the whole brain, as well as frontal, central, temporal, and posterior regions.

The developmental trajectory for each slope measure was plotted and analyzed quantitatively using Individual Growth Curve (IGC) models. Separate models were fit with slope or offset as the dependent variable (whole brain, frontal, central, temporal, posterior) and time point as the fixed effect. All models were fit with random intercepts and random slopes specified at each time point to account for variation at the individual level. To quantify changes between each time point, pairwise comparisons with Benjamini-Hochberg (BH) corrections were applied to correct for multiple comparisons. To investigate potential sex differences in developmental trajectories of slope, IGC models were re-fit with sex added as a factorial fixed effect, nested by time. If significant, sex was added as a covariate in subsequent models.

To investigate the potential associations between maternal anxiety symptoms and whole brain slope and offset over time, Linear Mixed Models (LMMs) were used. The maternal anxiety variables were transformed using the Box-Cox method to account for positive/right skew in the data, and subsequently standardized to facilitate interpretability. The models were specified with maternal anxiety symptoms, nested by time as the fixed effect, whole brain slope or offset as the outcome, and a random intercept specified. We focused on whole brain slope and offset for these analyses as they were exploratory and to limit the likelihood of encountering type 1 errors when running a large number of comparisons, or type 2 errors (if over-correcting for too many comparisons), and as prior research has primarily focused on whole brain slope.

## Results

### Sample Characteristics

Sociodemographic characteristics for the sample are presented in Table 1. Children in the sample were predominantly non-Hispanic White. Parental education levels and annual household income indicated that most families were middle to high socioeconomic status.

**Table 1.**
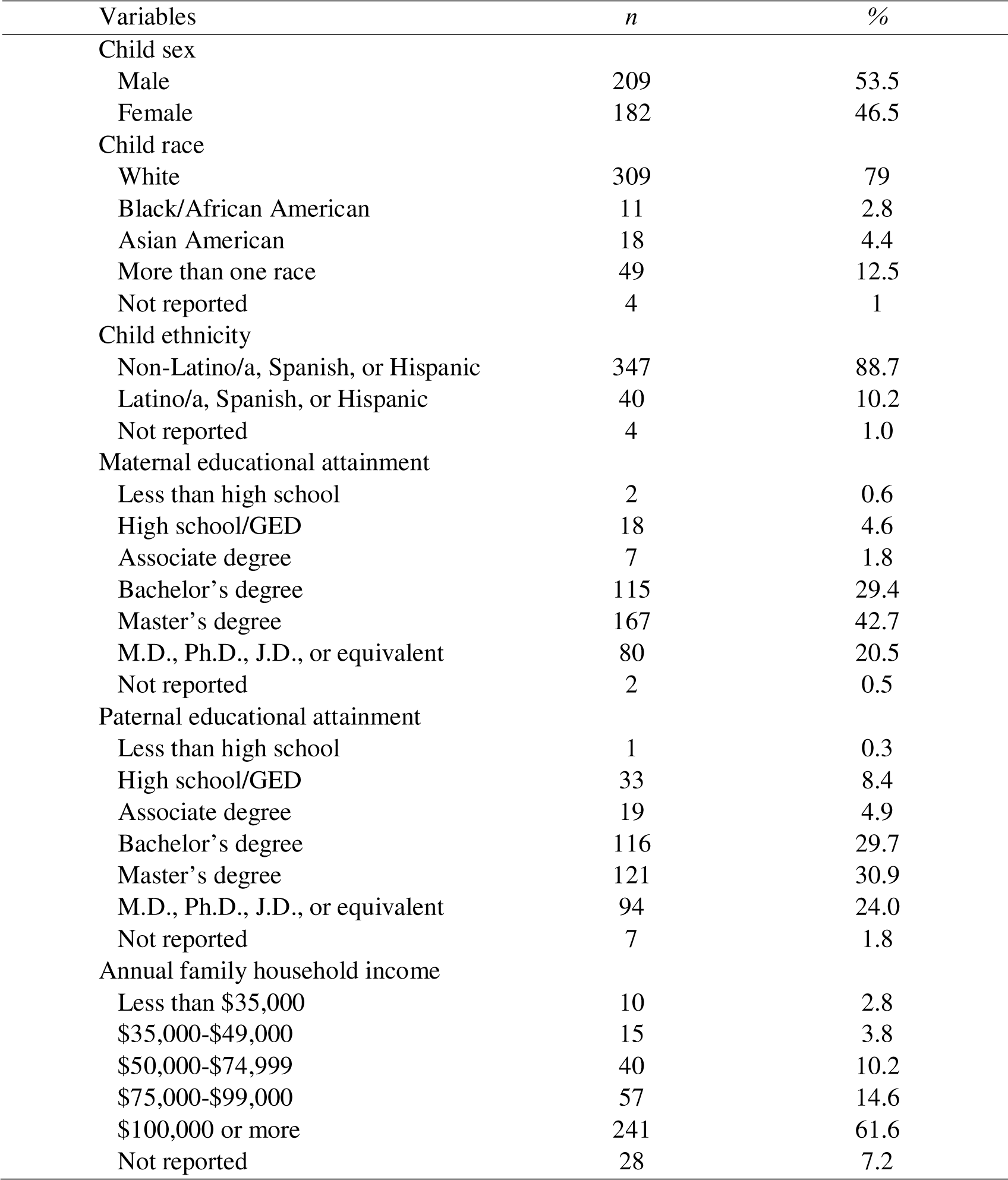
Parent and child demographic information collected at infancy (N=391).

| Variables | *n* | % |
| --- | --- | --- |
| Child sex |  |  |
| Male | 209 | 53.5 |
| Female | 182 | 46.5 |
| Child race |  |  |
| White | 309 | 79 |
| Black/African American | 11 | 2.8 |
| Asian American | 18 | 4.4 |
| More than one race | 49 | 12.5 |
| Not reported | 4 | 1 |
| Child ethnicity |  |  |
| Non-Latino/a, Spanish, or Hispanic | 347 | 88.7 |
| Latino/a, Spanish, or Hispanic | 40 | 10.2 |
| Not reported | 4 | 1.0 |
| Maternal educational attainment |  |  |
| Less than high school | 2 | 0.6 |
| High school/GED | 18 | 4.6 |
| Associate degree | 7 | 1.8 |
| Bachelor's degree | 115 | 29.4 |
| Master's degree | 167 | 42.7 |
| M.D., Ph.D., J.D., or equivalent | 80 | 20.5 |
| Not reported | 2 | 0.5 |
| Paternal educational attainment |  |  |
| Less than high school | 1 | 0.3 |
| High school/GED | 33 | 8.4 |
| Associate degree | 19 | 4.9 |
| Bachelor's degree | 116 | 29.7 |
| Master's degree | 121 | 30.9 |
| M.D., Ph.D., J.D., or equivalent | 94 | 24.0 |
| Not reported | 7 | 1.8 |
| Annual family household income |  |  |
| Less than \$35,000 | 10 | 2.8 |
| \$35,000-\$49,000 | 15 | 3.8 |
| \$50,000-\$74,999 | 40 | 10.2 |
| \$75,000-\$99,000 | 57 | 14.6 |
| \$100,000 or more | 241 | 61.6 |
| Not reported | 28 | 7.2 |

### Developmental Trajectories

The aperiodic fit and developmental trajectories of slope and offset for the whole brain at each time point are depicted in Figure 2. Longitudinal developmental trajectories for each of the aperiodic slope and offset variables were quantified using individual growth curve models, with Benjamini-Hochberg (BH) corrected pairwise comparisons between each age. Across the whole brain, slope increased significantly from infancy to 5 years (0.05, *p* <.001). There was a significant decrease from 5 to 7 years (-0.03, *p* = .039). Offset significantly increased from infancy to 3 years (0.10, *p* <.001), and then significantly decreased from 3 years to 7 years (-0.27, *p* <.001). Full model parameters and pairwise comparisons are provided in Supplementary Tables 1 and 2

**Figure 2.**
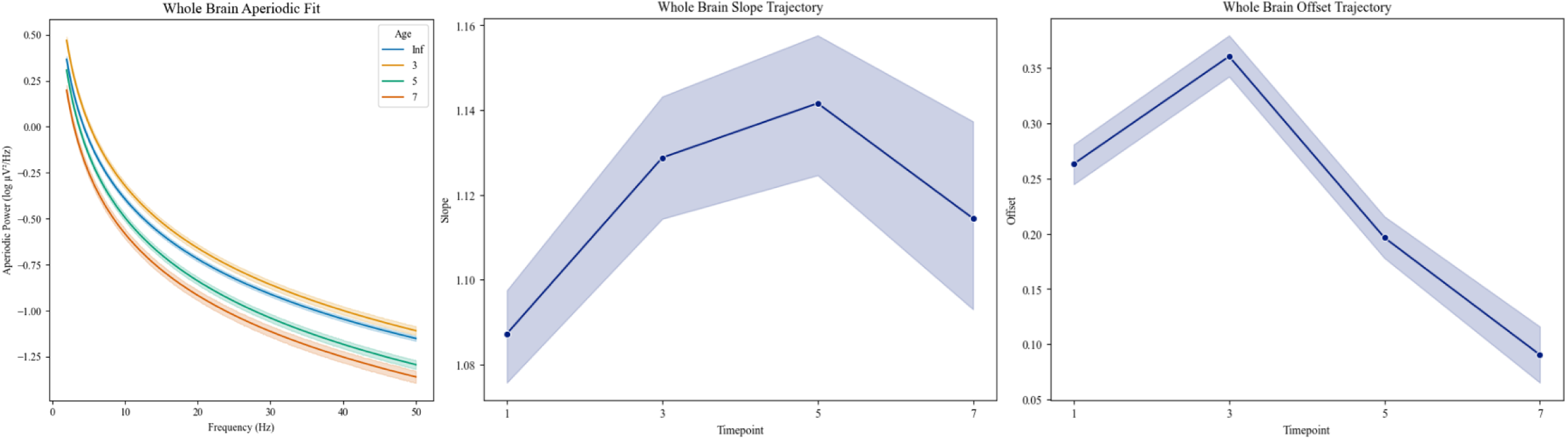
Aperiodic fit and slope and offset at each time point across whole brain.

#### Regional Differences

Changes in slope across each time point by region are depicted in Figure 3; aperiodic fit for each region is provided in Supplementary Figure 1. The pattern of developmental trajectories differed regionally. Across regions, significant increases were seen from infancy to 3 years (*ps* <.001). However, increases in frontal regions were decelerated through 5 years, whereas in temporal regions, they occurred at an accelerated rate. In central and posterior regions, significant decreases were observed from ages 3 years to 7 years (*ps* <.001). There were non-significant mean decreases in the frontal and temporal regions from 5 years to 7 years. Overall, slope began decreasing at age 3 years in posterior and central regions, but not until age 5 years in frontal and temporal regions. Offset increased from infancy to 3 years in frontal (*p* <.001), central (*p* = .004), and temporal (*p* <.001) regions, and then decreased to age 7 years (*ps* <.001). The greatest increase from infancy to 3 years was observed in the frontal region. Offset was stable from infancy to 3 years in the posterior region and then decreased to 7 years (*p* <.001). Full models and pairwise comparisons are provided in Supplementary Tables 3 to 13.

**Figure 3.**
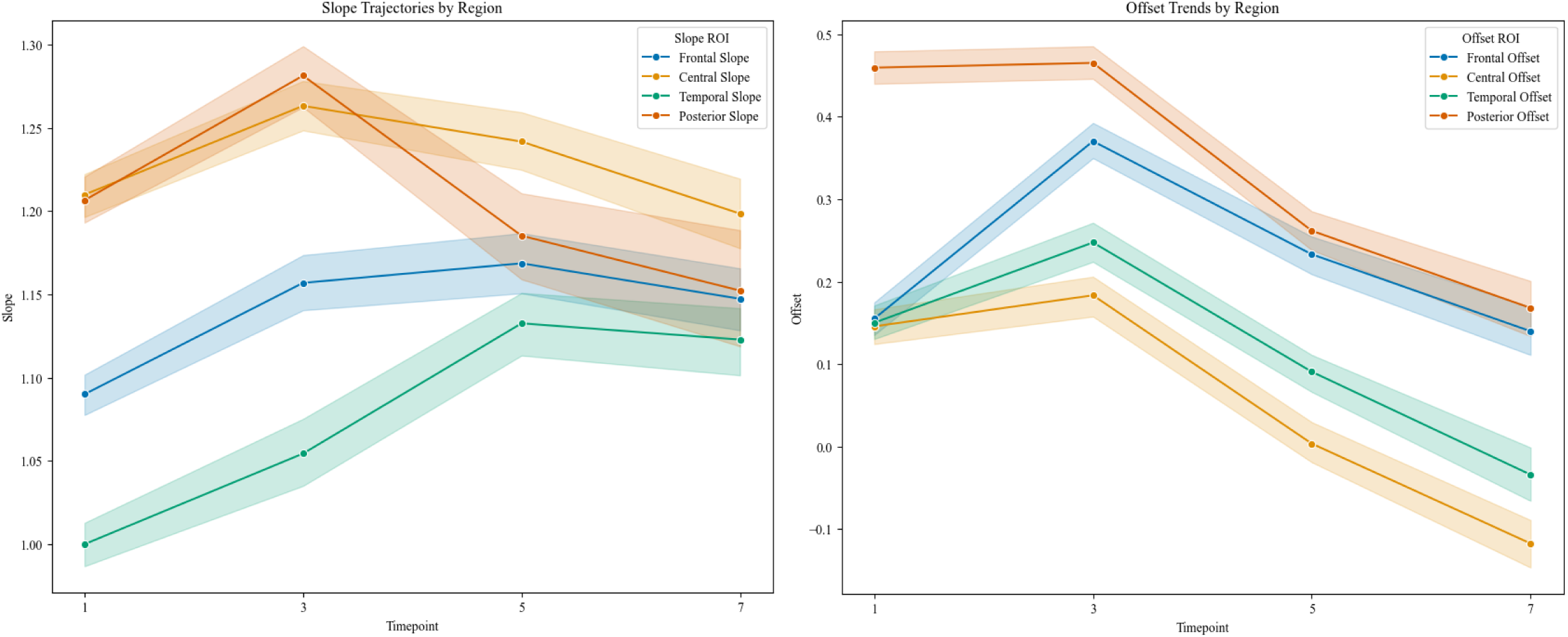
Developmental trajectories of slope and offset in frontal, central, temporal, and posterior ROIs.

#### Sex Differences

Child sex, nested by time, was added to the models to investigate potential sex differences. There were significant sex differences for slope across the whole brain. Female participants had steeper slope at 3 years (*b* = -0.04, *p*<.001) and 5 years (*b* = -0.07, *p* <.001). For offset across the whole brain, there was a significant sex difference, whereby offset was greater in males at 3 years (*b* = -0.38, *p* <.035). Trajectories for whole brain slope and offset stratified by sex are shown in Figure 4. There also were sex differences in slope in frontal and temporal regions and in offset in central and temporal regions. Slope was steeper in female participants in the frontal region at age 5 years (*b*=-0.06, *p*=.004), and in the temporal region at ages 3 years (*b*=-0.07, *p*<.001) and 5 years (*b*=-0.06, *p*=.001) (Supplementary Figure 2). Offset was greater in female participants in the central region at age 3 years (b=-0.54, p=.020), and in the temporal region at ages 3 years (b=-0.8, p<.001), 5 years (b=-0.11, p=.001, and 7 years (b=-0.8, p=.013) (Supplementary Figure 3).

**Figure 4.**
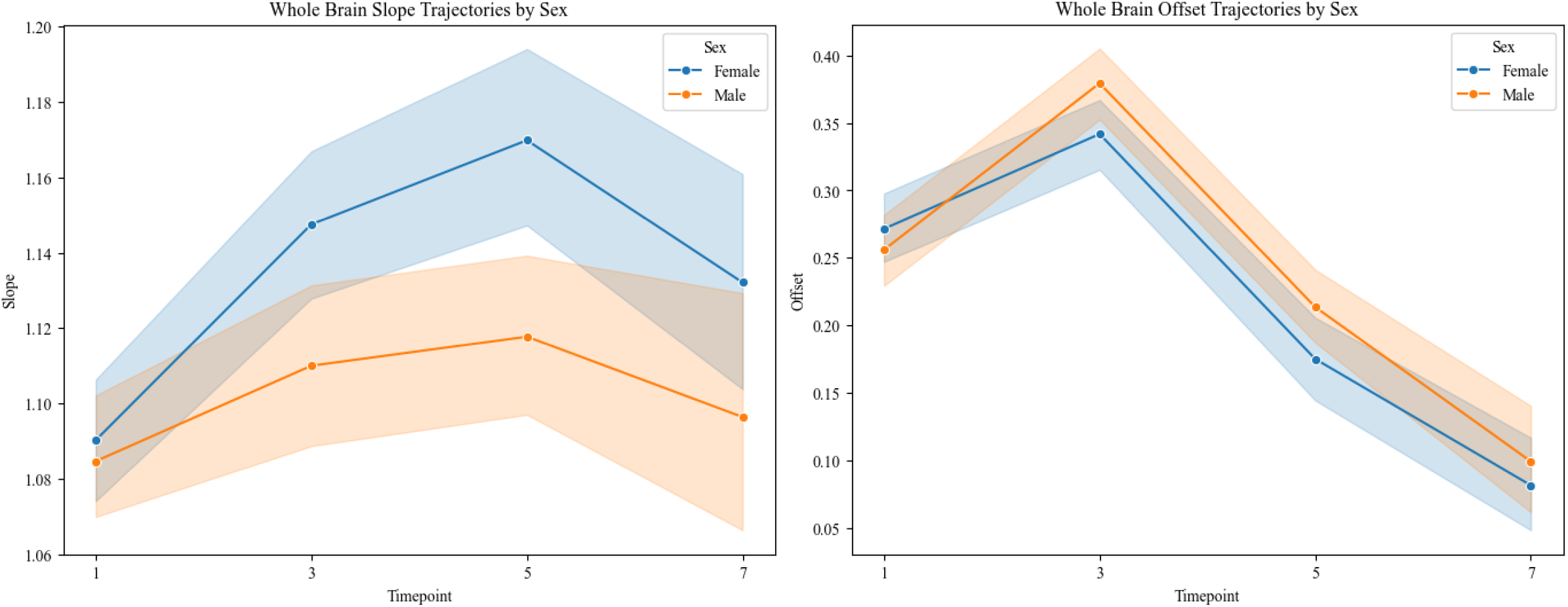
Developmental trajectories for aperiodic slope and offset stratified by sex in whole brain.

**Figure 5.**
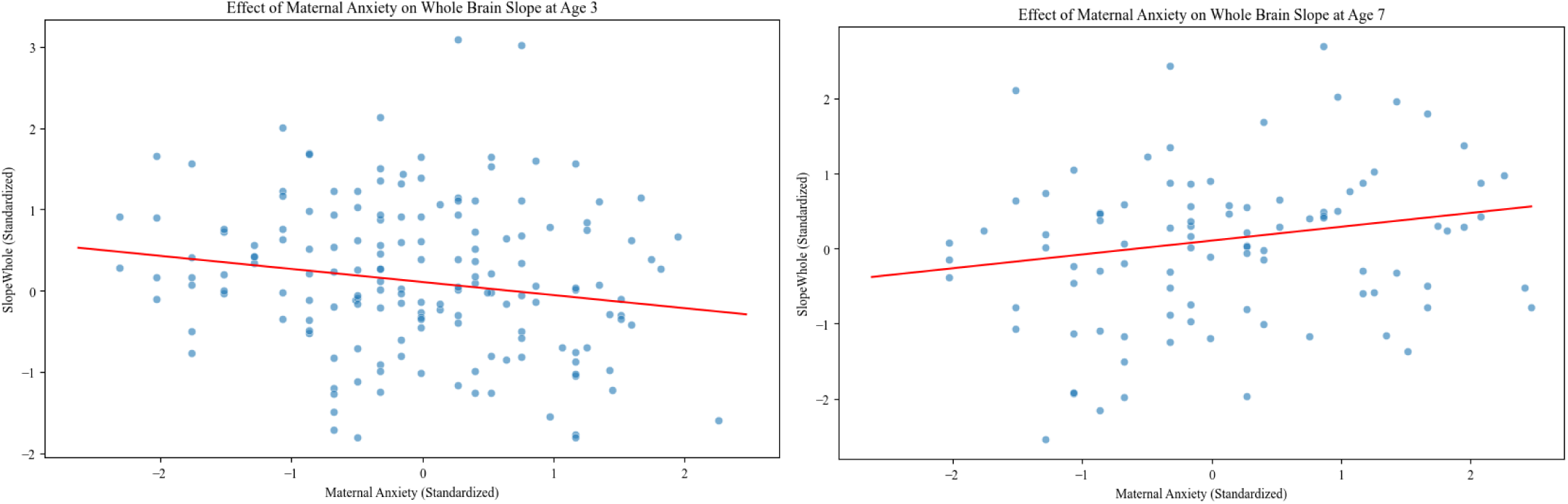
Scatterplots showing the significant associations of maternal anxiety with whole brain slope at 3 years and 7 years, respectively.

### Whole Brain Slope and Maternal Anxiety Symptoms

We investigated how maternal anxiety may be associated with slope and offset at each age using mixed effects models, with anxiety nested in each timepoint as the predictor and slope or offset as the outcome variable. The direction of observed associations varied by timepoint. There was a significant negative association between maternal anxiety symptoms and child slope at age 3 years, and a significant positive association between maternal anxiety symptoms and child slope at age 7 years (Table 3; Figure 8). There were no significant associations between maternal anxiety symptoms and offset.

**Table 2.** Descriptive statistics for the main study variables.

|  | Mean | SD | Min | Max |
| --- | --- | --- | --- | --- |
| Maternal anxiety symptoms (infancy) | 33.45 | 7.17 | 20 | 56 |
| Maternal anxiety symptoms (3 years) | 33.34 | 7.0 | 21 | 57 |
| Maternal anxiety symptoms (5 years) | 33.09 | 7.95 | 20 | 58 |
| Maternal anxiety symptoms (7 years) | 34.5 | 8.76 | 21 | 68 |

*Note.* Maternal anxiety symptoms were calculated from the trait anxiety form of the Spielberger State-Trait Anxiety Inventory (STAI-T).

**Table 3.** Mixed effects model for the associations between maternal anxiety symptoms, nested within time point, and EEG slope in the whole brain.

|  | Estimate | Std. Error | t | Sig. | 95% Confidence Interval |  |
| --- | --- | --- | --- | --- | --- | --- |
|  |  |  |  |  | Lower Bound | Upper Bound |
| Anxiety TP1 | 0.02 | 0.06 | 0.36 | .716 | -0.01 | 0.22 |
| **Anxiety TP3** | -0.16 | 0.08 | -2.08 | **.038\*** | -0.31 | -0.01 |
| Anxiety TP5 | 0.01 | 0.09 | 0.10 | .920 | -0.16 | 0.18 |
| **Anxiety TP7** | 0.19 | 0.09 | 2.06 | **.040\*** | 0.01 | 0.36 |
| **Sex (Male)** | -0.24 | 0.09 | -2.87 | **.004\*\*** | -0.40 | -0.08 |
| Participant Var | 0.15 |  |  |  |  |  |

**Table 4.** Mixed effects model for the associations between maternal anxiety symptoms, nested within time point, and EEG offset in the whole brain.

|  | Estimate | Std. Error | t | Sig. | 95% Confidence Interval |  |
| --- | --- | --- | --- | --- | --- | --- |
|  |  |  |  |  | Lower Bound | Upper Bound |
| Anxiety TP1 | 0.06 | 0.06 | 1.00 | .315 | -0.06 | 0.17 |
| Anxiety TP3 | -0.05 | 0.08 | -0.60 | .548 | -0.20 | 0.11 |
| Anxiety TP5 | 0.07 | 0.08 | 0.87 | .385 | -0.09 | .22 |
| Anxiety TP7 | -0.13 | 0.09 | -1.28 | .201 | 0.30 | 0.06 |
| Sex (Male) | 0.04 | 0.08 | 0.51 | .610 | -0.12 | 0.20 |
| Participant Var | 0.129 |  |  |  |  |  |

## Discussion

In the current study, we aimed to characterize the developmental trajectories of EEG aperiodic activity (slope and offset) from early to middle childhood in a longitudinal sample assessed from infancy (5-12 months) through 7 years of age. We investigated potential regional and sex differences. Further, we examined whether the nature of the association between child aperiodic activity and maternal anxiety symptoms varied by age, in line with changes in developmental trajectories. We observed nonlinear trajectories of the slope, with a pattern of increasing (steepening) slope during early childhood and decreasing (flattening) slope in middle childhood. These patterns of trajectories demonstrated differences by brain region and by sex. Additionally, the direction of associations between slope and maternal anxiety varied by age of assessment, in line with observed changes in the direction of the developmental trajectory.

### Developmental Trajectories

Longitudinally, there were distinct nonlinear developmental trajectories in slope and offset, which varied across brain regions. In the whole brain, slope increased from infancy to 5 years, with the greatest change from infancy to 3 years. There was a significant decrease from 5 to 7 years. Offset increased until 3 years, then decreased until 7 years. These results are consistent with those of McSweeney et al. (2023), who reported a nonlinear developmental trend in aperiodic activity (suggesting a quadratic model was a better fit compared to linear) in their cross-sectional sample of 4- to 11-year-old children. This transition from increasing to decreasing slope trajectories in whole brain regions may align with the shift Cellier et al. (2021) reported in dominant oscillatory activity from theta to alpha around 7 years. The observed trajectories also align with the complex systems framework, which posits that biological systems are characterized by nonlinear dynamics (Cohen et al., 2022).

Similar nonlinear trajectories have been observed in studies of developmental trajectories of cortical structure (Shaw et al., 2008). Consistent with the hypothesis that offset reflects broadband neuronal firing, the observed shift in trajectory may mark the broad transition from rapid neurodevelopmental growth that occurs in early childhood to synaptic pruning and ongoing systems optimization. Aperiodic slope is hypothesized to index E-I balance. Early childhood is characterized by unique maturational processes in inhibitory networks, including ongoing migration, maturation, and integration of GABAergic inhibitory neurons into the cortices that likely contribute to the early reduction in E-I ratio (Paredes et al., 2016; Wilkinson et al., 2024). As children continue to mature, ongoing network optimization may modulate the GABAergic and glutamatergic “push-pull” mechanism, resulting in a shift towards increased excitatory tone (Hill et al., 2022; McKeon et al., 2024).

In analyses that examined trajectories by region, slope increased significantly from infancy through age 5 years, before stabilizing from 5 to 7 years of age, in the frontal and temporal regions. In the central and posterior regions, we found significant decreases in slope earlier, from 3 to 7 years. While offset increased from infancy to 3 years and then decreased to 7 years in frontal, central, and temporal regions, the greatest increase from infancy to 3 years was seen in the frontal region; offset remained stable in the posterior region from infancy to 3 years. Cellier et al. (2021) investigated age-related trends in aperiodic activity using linear regression modeling in two independent samples between 3 to 24 years of age. They observed a significant negative association between age and both anterior and posterior offset starting from 3 years, but only posterior for slope. In the present study, there were decreases from 3 years in posterior slope, posterior offset, and anterior offset, whereas frontal slope continued to increase up to 5 years, with relative stabilization until 7 years. These results likely correspond with the null results in anterior slope observed by Cellier et al. (2021) when adopting a linear approach.

Our findings highlight differential regional development in aperiodic activity, which may reflect the caudal–rostral pattern of development observed in the cortex (e.g., Shaw et al. (2008). Cortical development follows a tightly regulated genetic ‘blueprint’ (Silbereis et al., 2016). Accelerated development in more posterior regions may culminate in an earlier shift from increasing to decreasing aperiodic activity. Importantly, slope and offset are inherently correlated; prior studies with cross-sectional, linear designs have reported similar trends (i.e., a linear decrease) for both during aging. The unique trajectories of slope and offset in this study provide further support for the notion that they index unique neurological processes (i.e., E-I balance vs. overall neuronal population spiking, respectively). The relative “lag” observed in frontal slope may reflect extended maturation of E-I related processes in the frontal cortex, which is necessary for the development of higher-order cognitive functions.

We identified significant sex differences, whereby females exhibited greater increases in (steeper) slope relative to males. These differences were most pronounced at 3 years and 5 years in the whole brain, frontal, and temporal regions. To a lesser degree, sex differences were also seen in offset in the whole brain, central, and temporal regions. Wilkinson et al. (2024) also reported significant sex effects in aperiodic activity in their study from infancy to 3 years. Sex differences appeared more pronounced at 3 years and 5 years in the present study. Sex-based differences in brain development during childhood are well established. For example, sex effects have been reported in studies of EEG power, functional connectivity, and microstates (Bagdasarov et al., 2022; Gartstein et al., 2020; Kavčič et al., 2023). McSweeney et al. (2021) observed significant sex differences in aperiodic activity in adolescence. Thus, results across these studies suggest sex effects in the EEG slope are apparent after infancy and extend through at least adolescence. Overall, childhood appears to be a dynamic period with significant development in brain function, including age-by-sex interactions.

### Associations of Whole-Brain Aperiodic Activity with Maternal Anxiety Symptoms from Infancy to 7 Years

We tested longitudinal associations between whole brain slope and offset with maternal anxiety from infancy to age 7 years. We observed significant associations between maternal anxiety symptoms and child slope at ages 3 years and 7 years. The nature of the associations differed: At 3 years, the association was negative, whereas at 7 years, the association was positive. The direction of associations corresponded with increasing slope trajectories at 3 years and an emerging pattern of decreasing slope trajectories at 7 years. Importantly, rather than demonstrating a consistent direction of association (e.g., greater maternal anxiety always associated with flatter slope), maternal anxiety appeared to be associated with dynamic changes in developmental trajectory for slope. In other words, maternal anxiety symptoms were differentially associated with slope at different ages (i.e., negative versus positive). This is an important consideration for future research, as these results suggest that the interpretation of potential implications of exposure to maternal psychopathology on child brain development, as reflected in measures of slope, must consider the timing of assessment.

### Strengths and Limitations

Strengths of this study include the large longitudinal sample, multimodal measures, and investigation of timing effects. The findings should be considered within the context of the study’s limitations. Aperiodic EEG measures are likely susceptible to non-neurophysiological developmental changes in anatomy that occur during early childhood, such as increases in skull thickness, volume of cerebrospinal fluid, and closure of the anterior fontanel, all of which are difficult to measure and therefore control for in analyses. We focused on aperiodic EEG activity given emerging interest in this area and the need for longitudinal analyses of developmental trajectories. Thus, we did not consider periodic measures. Further research is required to determine what the slope indexes biologically, with the E-I hypothesis based primarily on theoretical models and animal research. Additional research is also needed to orient the current findings in the context of periodic EEG. The current sample comprised families who are predominantly White and of middle to high socioeconomic status, which may limit the generalizability of the findings. Finally, although a strength of this study included its longitudinal design, the 2-year gaps between assessments limited our ability to identify nuanced age-related trends and transition points in nonlinear slope and offset development.

## Conclusions

Our longitudinal findings indicate nonlinear trajectories in EEG aperiodic activity during childhood, which varied by brain region and by sex. The results emphasize the dynamic nature of cortical development in childhood. Differential developmental trajectories of aperiodic activity are likely underpinned by regionally differential timing in shifts from rapid early brain growth to network optimization and synaptic pruning. The findings add to a growing body of evidence that slope and offset index unique neurological processes (i.e., E-I balance vs. overall neuronal population spiking, respectively). The changing nature of the associations between maternal anxiety and slope across ages emphasizes the importance of considering how normative developmental changes in slope trajectories may influence associations with putative risk factors in future research. These considerations will be crucial, both when investigating the biological mechanisms that slope and offset index and when investigating associations with other important developmental factors.

