## Supplement for "Longitudinal trajectories of aperiodic EEG activity in early to middle childhood"

Supplementary Figure 1

*Aperiodic fit for frontal, central, temporal, and posterior regions at each time point*

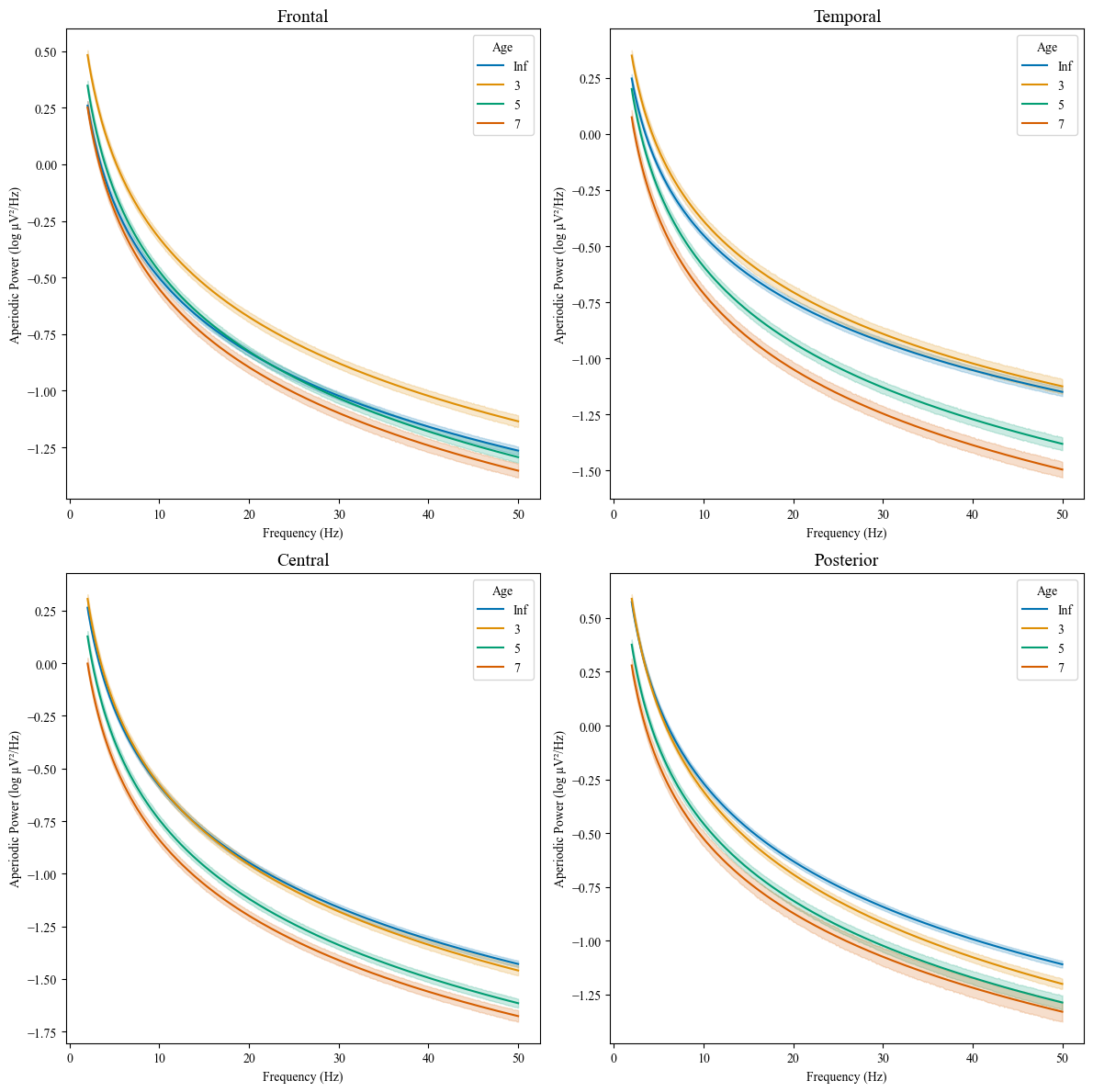

Supplementary Figure 2

*Developmental trajectories for aperiodic slope stratified by sex in frontal, temporal, and temporal regions.*

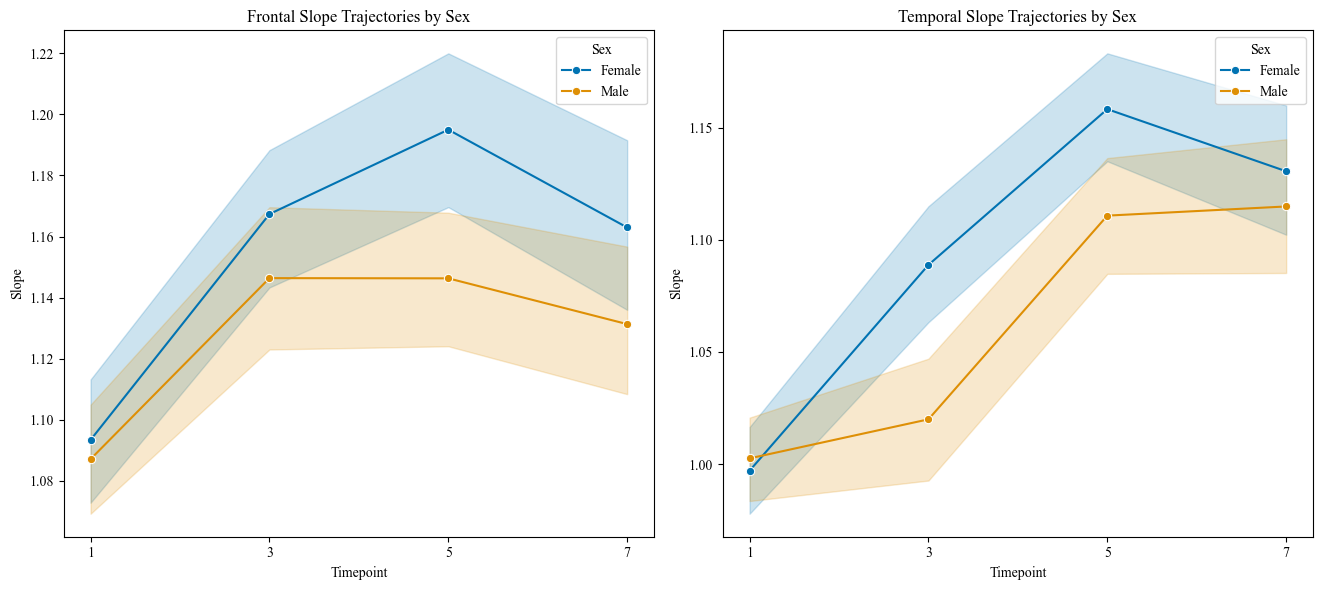

*Slope was greater in female participants in the frontal region at age 5 years (b=-0.06, p=.004), and in the temporal region at ages 3 years (b=-0.07, p<.001) and 5 years (b=-0.06, p=.001).*

Supplementary Figure 3

*Developmental trajectories for aperiodic offset stratified by sex in central, and temporal regions*

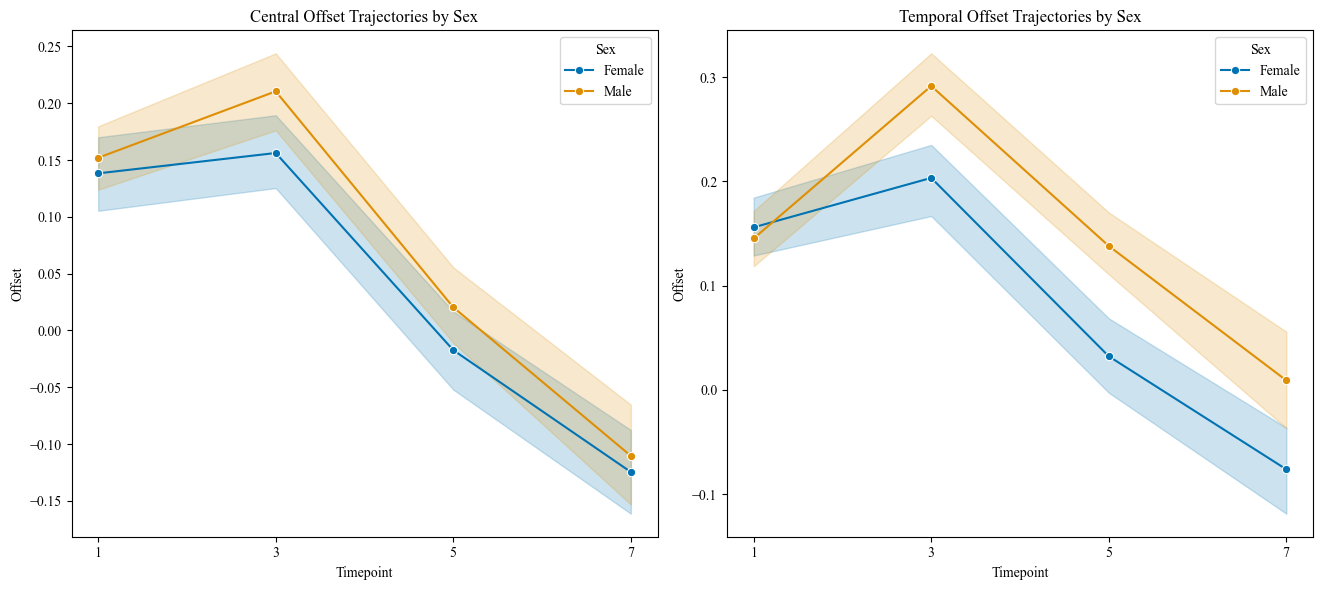

*Offset was greater in female participants in the central region at age 3 years (b=-0.54, p=.020), and in the temporal region at ages 3 years (b=-0.8, p<.001), 5 years (b=-0.11, p=.001, and 7 years (b=-0.8, p=.013).*

### Supplementary Table 1 (A)

### Individual growth curve/mixed effects model for whole brain slope from infancy to 7 years.

|  | | | | | 95% Confidence Interval | |
| --- | --- | --- | --- | --- | --- | --- |
|  | Estimate | Std. Error | t | Sig. | Lower Bound | Upper Bound |
| Intercept (infancy) | 1.09 | 0.01 | 186.71 | <.001 | 1.08 | 1.10 |
| 3 years | 0.04 | 0.01 | 4.51 | <.001 | 0.02 | 0.06 |
| 5 years | 0.05 | 0.01 | 5.29 | <.001 | 0.03 | 0.07 |
| 7 years | 0.03 | 0.01 | 2.10 | .036 | 0.00 | 0.05 |
| Group Var | 0.01 |  |  |  |  |  |
| Group x Age 3 Cov | 0.00 |  |  |  |  |  |
| Age 3 Var | 0.01 |  |  |  |  |  |
| Group x Age 5 Cov | 0.00 |  |  |  |  |  |
| Age 3 x Age 5 Cov | 0.01 |  |  |  |  |  |
| Age 5 Var | 0.01 |  |  |  |  |  |
| Group x Age 7 Cov | 0.00 |  |  |  |  |  |
| Age 3 x Age 7 Cov | 0.01 |  |  |  |  |  |
| Age 5 x Age 7 Cov | 0.01 |  |  |  |  |  |
| Age 7 Var | 0.01 |  |  |  |  |  |

### Supplementary Table 1 (B)

### Pairwise Comparisons

|  |  | Diff | Sig. |
| --- | --- | --- | --- |
| 3 | 5 | 0.01 | .282 |
|  | 7 | -0.02 | .195 |
| 5 | 7 | -0.03 | .039 |

### Supplementary Figure 4 (C)

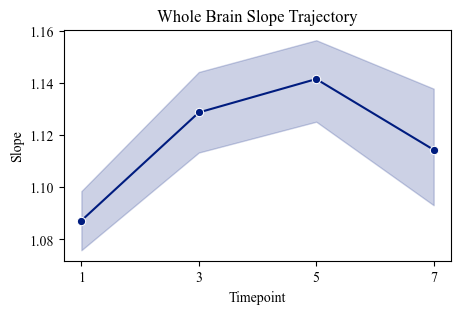

### Whole Brain Slope Trajectory

### Supplementary Table 2 (A)

### Individual growth curve/mixed effects model for frontal slope from infancy to 7 years.

|  | | | | | 95% Confidence Interval | |
| --- | --- | --- | --- | --- | --- | --- |
|  | Estimate | Std. Error | t | Sig. | Lower Bound | Upper Bound |
| Intercept (infancy) | 1.09 | 0.01 | 165.69 | <.001 | 1.08 | 1.10 |
| 3 years | 0.07 | 0.01 | 6.51 | <.001 | 0.05 | 0.09 |
| 5 years | 0.08 | 0.01 | 7.23 | <.001 | 0.06 | 0.10 |
| 7 years | 0.06 | 0.01 | 4.83 | <.001 | 0.03 | 0.08 |
| Group Var | 0.01 |  |  |  |  |  |
| Group x Age 3 Cov | -0.01 |  |  |  |  |  |
| Age 3 Var | 0.01 |  |  |  |  |  |
| Group x Age 5 Cov | -0.01 |  |  |  |  |  |
| Age 3 x Age 5 Cov | 0.01 |  |  |  |  |  |
| Age 5 Var | 0.01 |  |  |  |  |  |
| Group x Age 7 Cov | -0.01 |  |  |  |  |  |
| Age 3 x Age 7 Cov | 0.01 |  |  |  |  |  |
| Age 5 x Age 7 Cov | 0.01 |  |  |  |  |  |
| Age 7 Var | 0.01 |  |  |  |  |  |

### Supplementary Table 2 (B)

### Pairwise Comparisons

|  |  | Diff | Sig. |
| --- | --- | --- | --- |
| 3 | 5 | 0.01 | .344 |
|  | 7 | -0.01 | .409 |
| 5 | 7 | -0.02 | .098 |

### Supplementary Figure 5 (C)

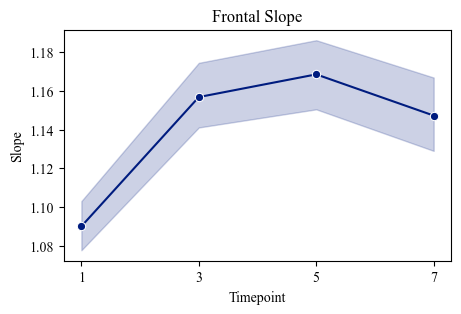

### Frontal Slope Trajectory

### Supplementary Table 3 (A)

### Individual growth curve/mixed effects model for central slope from infancy to 7 years.

|  | | | | | 95% Confidence Interval | |
| --- | --- | --- | --- | --- | --- | --- |
|  | Estimate | Std. Error | t | Sig. | Lower Bound | Upper Bound |
| Intercept (infancy) | 1.21 | 0.01 | 189.11 | <.001 | 1.20 | 1.22 |
| 3 years | 0.05 | 0.01 | 5.58 | <.001 | 0.04 | 0.07 |
| 5 years | 0.03 | 0.01 | 2.98 | .003 | 0.01 | 0.05 |
| 7 years | -0.01 | 0.01 | -0.90 | .367 | -0.03 | 0.01 |
| Group Var | 0.01 |  |  |  |  |  |
| Group x Age 3 Cov | -0.01 |  |  |  |  |  |
| Age 3 Var | 0.01 |  |  |  |  |  |
| Group x Age 5 Cov | -0.01 |  |  |  |  |  |
| Age 3 x Age 5 Cov | 0.01 |  |  |  |  |  |
| Age 5 Var | 0.01 |  |  |  |  |  |
| Group x Age 7 Cov | -0.01 |  |  |  |  |  |
| Age 3 x Age 7 Cov | 0.01 |  |  |  |  |  |
| Age 5 x Age 7 Cov | 0.01 |  |  |  |  |  |
| Age 7 Var | 0.01 |  |  |  |  |  |

### Supplementary Table 3 (B)

### Pairwise Comparisons

|  |  | Diff | Sig. |
| --- | --- | --- | --- |
| 3 | 5 | -0.02 | .030 |
|  | 7 | -0.06 | <.001 |
| 5 | 7 | -0.04 | .001 |

### Supplementary Figure 6 (C)

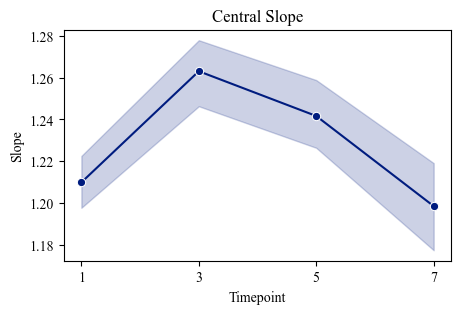

### Central Slope Trajectory

### Supplementary Table 4 (A)

|  | | | | | 95% Confidence Interval | |
| --- | --- | --- | --- | --- | --- | --- |
|  | Estimate | Std. Error | t | Sig. | Lower Bound | Upper Bound |
| Intercept (infancy) | 1.00 | 0.01 | 146.02 | <.001 | 0.99 | 1.01 |
| 3 years | 0.05 | 0.01 | 4.60 | <.001 | 0.03 | 0.08 |
| 5 years | 0.13 | 0.01 | 12.23 | <.001 | 0.11 | 0.15 |
| 7 years | 0.12 | 0.01 | 9.92 | <.001 | 0.09 | 0.14 |
| Group Var | 0.01 |  |  |  |  |  |
| Group x Age 3 Cov | -0.01 |  |  |  |  |  |
| Age 3 Var | 0.02 |  |  |  |  |  |
| Group x Age 5 Cov | -0.01 |  |  |  |  |  |
| Age 3 x Age 5 Cov | 0.01 |  |  |  |  |  |
| Age 5 Var | 0.01 |  |  |  |  |  |
| Group x Age 7 Cov | -0.01 |  |  |  |  |  |
| Age 3 x Age 7 Cov | 0.01 |  |  |  |  |  |
| Age 5 x Age 7 Cov | 0.01 |  |  |  |  |  |
| Age 7 Var | 0.01 |  |  |  |  |  |

### Individual growth curve/mixed effects model for temporal slope from infancy to 7 years.

### Supplementary Table 4 (B)

### Pairwise Comparisons

|  |  | Diff | Sig. |
| --- | --- | --- | --- |
| 3 | 5 | 0.08 | <.001 |
|  | 7 | 0.06 | <.001 |
| 5 | 7 | -0.02 | 0.223 |

### Supplementary Figure 7(C)

### Temporal Slope Trajectory

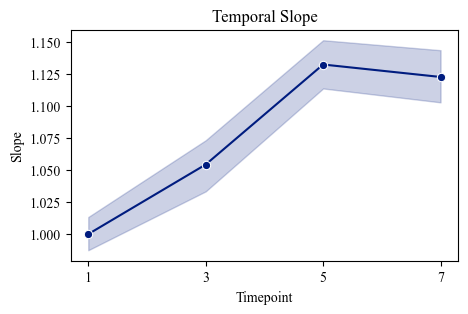

### Supplementary Table 5 (A)

### Individual growth curve/mixed effects model for posterior offset from infancy to 7 years.

|  | | | | | 95% Confidence Interval | |
| --- | --- | --- | --- | --- | --- | --- |
|  | Estimate | Std. Error | t | Sig. | Lower Bound | Upper Bound |
| Intercept (infancy) | 1.21 | 0.01 | 167.68 | <.001 | 1.19 | 1.22 |
| 3 years | 0.08 | 0.01 | 6.34 | <.001 | 0.05 | 0.10 |
| 5 years | -0.03 | 0.01 | -1.88 | .060 | -0.06 | 0.00 |
| 7 years | -0.05 | 0.02 | -2.79 | .005 | -0.09 | -0.02 |
| Group Var | 0.00 |  |  |  |  |  |
| Group x Age 3 Cov | 0.00 |  |  |  |  |  |
| Age 3 Var | 0.01 |  |  |  |  |  |
| Group x Age 5 Cov | 0.00 |  |  |  |  |  |
| Age 3 x Age 5 Cov | 0.01 |  |  |  |  |  |
| Age 5 Var | 0.01 |  |  |  |  |  |
| Group x Age 7 Cov | 0.00 |  |  |  |  |  |
| Age 3 x Age 7 Cov | 0.01 |  |  |  |  |  |
| Age 5 x Age 7 Cov | 0.01 |  |  |  |  |  |
| Age 7 Var | 0.02 |  |  |  |  |  |

### Supplementary Table 5 (B)

### Pairwise Comparisons

|  |  | Diff | Sig. |
| --- | --- | --- | --- |
| 3 | 5 | -0.10 | <.001 |
|  | 7 | -0.13 | <.001 |
| 5 | 7 | -0.02 | 0.240 |

### Supplementary Figure 8 (C)

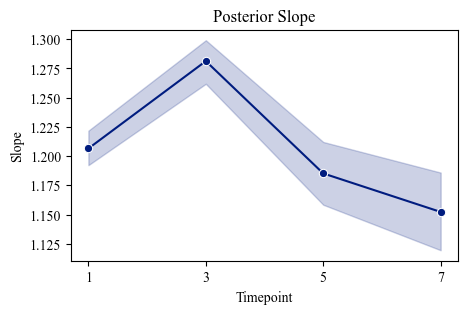

### Posterior Slope Trajectory

### Supplementary Table 6 (A)

### Individual growth curve/mixed effects model for whole brain offset from infancy to 7 years.

|  | | | | | 95% Confidence Interval | |
| --- | --- | --- | --- | --- | --- | --- |
|  | Estimate | Std. Error | t | Sig. | Lower Bound | Upper Bound |
| Intercept (infancy) | 0.26 | 0.01 | 29.00 | <.001 | 0.25 | 0.28 |
| 3 years | 0.10 | 0.01 | 8.64 | <.001 | 0.08 | 0.12 |
| 5 years | -0.07 | 0.01 | -5.47 | <.001 | -0.09 | -0.04 |
| 7 years | -0.17 | 0.01 | -12.16 | <.001 | -0.20 | -0.14 |
| Group Var | 0.02 |  |  |  |  |  |
| Group x Age 3 Cov | -0.01 |  |  |  |  |  |
| Age 3 Var | 0.01 |  |  |  |  |  |
| Group x Age 5 Cov | -0.02 |  |  |  |  |  |
| Age 3 x Age 5 Cov | 0.02 |  |  |  |  |  |
| Age 5 Var | 0.02 |  |  |  |  |  |
| Group x Age 7 Cov | -0.01 |  |  |  |  |  |
| Age 3 x Age 7 Cov | 0.01 |  |  |  |  |  |
| Age 5 x Age 7 Cov | 0.01 |  |  |  |  |  |
| Age 7 Var | 0.01 |  |  |  |  |  |

### Supplementary Table 6 (B)

### Pairwise Comparisons

|  |  | Diff | Sig. |
| --- | --- | --- | --- |
| 3 | 5 | -0.17 | <.001 |
|  | 7 | -0.27 | <.001 |
| 5 | 7 | -0.10 | <.001 |

### Supplementary Figure 9 (C)

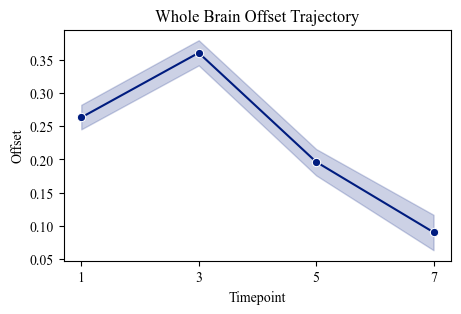

### Whole Brain Offset Trajectory

### Supplementary Table 7 (A)

### Individual growth curve/mixed effects model for frontal offset from infancy to 7 years.

|  | | | | | 95% Confidence Interval | |
| --- | --- | --- | --- | --- | --- | --- |
|  | Estimate | Std. Error | t | Sig. | Lower Bound | Upper Bound |
| Intercept (infancy) | 0.16 | 0.01 | 14.54 | <.001 | 0.13 | 0.18 |
| 3 years | 0.22 | 0.01 | 15.64 | <.001 | 0.19 | 0.24 |
| 5 years | 0.08 | 0.01 | 5.36 | <.001 | 0.05 | 0.11 |
| 7 years | -0.01 | 0.02 | -0.57 | .570 | -0.04 | 0.02 |
| Group Var | 0.03 |  |  |  |  |  |
| Group x Age 3 Cov | -0.02 |  |  |  |  |  |
| Age 3 Var | 0.03 |  |  |  |  |  |
| Group x Age 5 Cov | -0.02 |  |  |  |  |  |
| Age 3 x Age 5 Cov | 0.02 |  |  |  |  |  |
| Age 5 Var | 0.03 |  |  |  |  |  |
| Group x Age 7 Cov | -0.02 |  |  |  |  |  |
| Age 3 x Age 7 Cov | 0.02 |  |  |  |  |  |
| Age 5 x Age 7 Cov | 0.03 |  |  |  |  |  |
| Age 7 Var | 0.03 |  |  |  |  |  |

### Supplementary Table 7 (B)

### Pairwise Comparisons

|  |  | Diff | Sig. |
| --- | --- | --- | --- |
| 3 | 5 | -0.14 | <.001 |
|  | 7 | -0.23 | <.001 |
| 5 | 7 | -0.09 | <.001 |

### Supplementary Figure 10 (C)

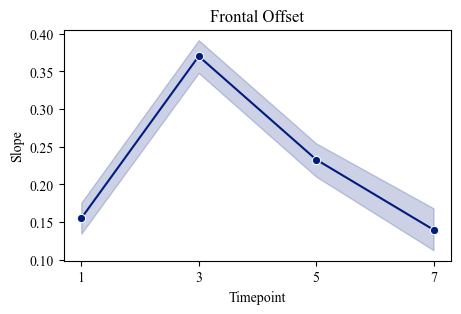

### Frontal Offset Trajectory

### Supplementary Table 8

### Individual growth curve/mixed effects model for central offset from infancy to 7 years.

|  | | | | | 95% Confidence Interval | |
| --- | --- | --- | --- | --- | --- | --- |
|  | Estimate | Std. Error | t | Sig. | Lower Bound | Upper Bound |
| Intercept (infancy) | 0.15 | 0.01 | 13.62 | <.001 | 0.13 | 0.17 |
| 3 years | 0.04 | 0.01 | 2.87 | .004 | 0.01 | 0.07 |
| 5 years | -0.15 | 0.02 | -9.49 | <.001 | -0.18 | -0.12 |
| 7 years | -0.26 | 0.02 | -15.60 | <.001 | -0.29 | -0.23 |
| Group Var | 0.03 |  |  |  |  |  |
| Group x Age 3 Cov | -0.02 |  |  |  |  |  |
| Age 3 Var | 0.02 |  |  |  |  |  |
| Group x Age 5 Cov | -0.02 |  |  |  |  |  |
| Age 3 x Age 5 Cov | 0.02 |  |  |  |  |  |
| Age 5 Var | 0.03 |  |  |  |  |  |
| Group x Age 7 Cov | -0.02 |  |  |  |  |  |
| Age 3 x Age 7 Cov | 0.02 |  |  |  |  |  |
| Age 5 x Age 7 Cov | 0.02 |  |  |  |  |  |
| Age 7 Var | 0.02 |  |  |  |  |  |

### Supplementary Table 8 (B)

### Pairwise Comparisons

|  |  | Diff | Sig. |
| --- | --- | --- | --- |
| 3 | 5 | -0.19 | <.001 |
|  | 7 | -0.30 | <.001 |
| 5 | 7 | -0.11 | <.001 |

### Supplementary Figure 11 (C)

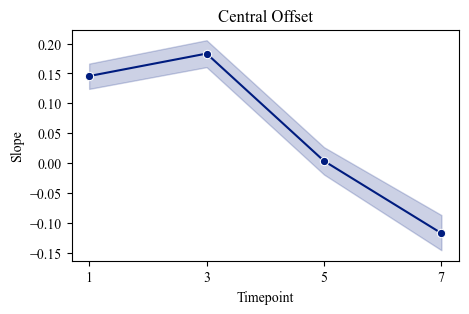

### Central Offset Trajectory

### Supplementary Table 9 (A)

|  | | | | | 95% Confidence Interval | |
| --- | --- | --- | --- | --- | --- | --- |
|  | Estimate | Std. Error | t | Sig. | Lower Bound | Upper Bound |
| Intercept (infancy) | 0.15 | 0.01 | 14.99 | <.001 | 0.13 | 0.17 |
| 3 years | 0.10 | 0.02 | 6.79 | <.001 | 0.07 | 0.13 |
| 5 years | -0.06 | 0.02 | -3.97 | <.001 | -0.09 | -0.03 |
| 7 years | -0.18 | 0.02 | -10.13 | <.001 | -0.22 | -0.15 |
| Group Var | 0.02 |  |  |  |  |  |
| Group x Age 3 Cov | -0.02 |  |  |  |  |  |
| Age 3 Var | 0.02 |  |  |  |  |  |
| Group x Age 5 Cov | -0.02 |  |  |  |  |  |
| Age 3 x Age 5 Cov | 0.02 |  |  |  |  |  |
| Age 5 Var | 0.02 |  |  |  |  |  |
| Group x Age 7 Cov | -0.02 |  |  |  |  |  |
| Age 3 x Age 7 Cov | 0.02 |  |  |  |  |  |
| Age 5 x Age 7 Cov | 0.02 |  |  |  |  |  |
| Age 7 Var | 0.02 |  |  |  |  |  |

### Individual growth curve/mixed effects model for temporal offset from infancy to 7 years.

### Supplementary Table 9 (B)

### Pairwise Comparisons

|  |  | Diff | Sig. |
| --- | --- | --- | --- |
| 3 | 5 | -0.16 | <.001 |
|  | 7 | -0.28 | <.001 |
| 5 | 7 | -0.12 | <.001 |

### Supplementary Figure 12 (C)

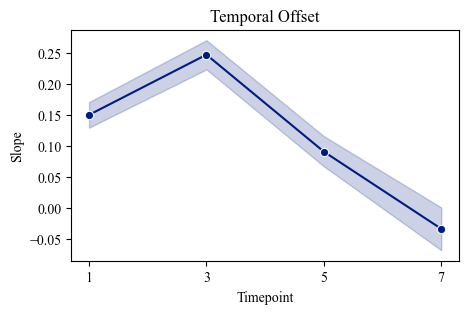

### Temporal Offset Trajectory

### Supplementary Table 10 (A)

### Individual growth curve/mixed effects model for posterior offset from infancy to 7 years.

|  | | | | | 95% Confidence Interval | |
| --- | --- | --- | --- | --- | --- | --- |
|  | Estimate | Std. Error | t | Sig. | Lower Bound | Upper Bound |
| Intercept (infancy) | 0.46 | 0.01 | 45.46 | <.001 | 0.44 | 0.48 |
| 3 years | 0.01 | 0.01 | 0.79 | .432 | -0.02 | 0.04 |
| 5 years | -0.20 | 0.02 | -13.36 | <.001 | -0.23 | -0.17 |
| 7 years | -0.29 | 0.02 | -15.17 | <.001 | -0.32 | -0.25 |
| Group Var | 0.02 |  |  |  |  |  |
| Group x Age 3 Cov | -0.02 |  |  |  |  |  |
| Age 3 Var | 0.02 |  |  |  |  |  |
| Group x Age 5 Cov | -0.02 |  |  |  |  |  |
| Age 3 x Age 5 Cov | 0.02 |  |  |  |  |  |
| Age 5 Var | 0.03 |  |  |  |  |  |
| Group x Age 7 Cov | -0.02 |  |  |  |  |  |
| Age 3 x Age 7 Cov | 0.02 |  |  |  |  |  |
| Age 5 x Age 7 Cov | 0.02 |  |  |  |  |  |
| Age 7 Var | 0.03 |  |  |  |  |  |

### Supplementary Table 10 (B)

### Pairwise Comparisons

|  |  | Diff | Sig. |
| --- | --- | --- | --- |
| 3 | 5 | -0.21 | <.001 |
|  | 7 | -0.30 | <.001 |
| 5 | 7 | -0.09 | <.001 |

### Supplementary Figure 13 (C)

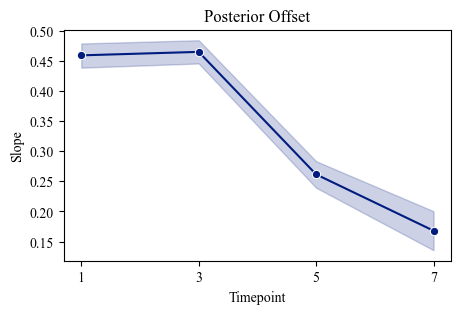

### Posterior Offset Trajectory
